# Kindlin-1 loss disrupts vascular and extracellular matrix organisation to sustain hypoxia in cutaneous squamous cell carcinoma

**DOI:** 10.64898/2026.06.01.728760

**Authors:** David Hardman, Giovana Carrasco, Martin Lee, Muhammad Furqan, Romain Enjalbert, Valerie G. Brunton, Miguel O. Bernabeu

## Abstract

Kindlin-1, encoded by *FERMT1*, is an essential integrin co-activator that regulates cell–extracellular matrix (ECM) adhesion, tissue architecture, and microenvironment signalling. Loss-of-function mutations in *FERMT1* cause Kindler epidermolysis bullosa, which is strongly associated with aggressive cutaneous squamous cell carcinoma (cSCC). Although Kindlin-1 deficiency promotes hypoxia and invasion, the impacts on ECM–vascular organisation and oxygen homeostasis are not known.

Here, using genetic deletion of Kindlin-1 in a murine model of cSCC across 2D cultures, 3D spheroids, and in vivo tumours, combined with collagen and vascular imaging and spatial mixed-effects modelling, we show that Kindlin-1 loss uncouples ECM–vascular regulation, driving hypoxia and tumour progression. Tumours in which Kindlin-1 was deleted displayed a dense but dysfunctional vascular network, with reduced tissue-to-vessel and inter-bifurcation distances, increased vessel alignment, and persistent hypoxia despite increased vascular density. Collagen deposition was reduced and fibres were straighter, indicating a simplified, invasion-permissive matrix. Hypoxia increased *Vegfa* and *Angpt1* expression while reducing *Col1a1*, and hypoxia-responsive spheroids confirmed greater hypoxia and invasiveness in Kindlin-1-deficient cells. Transcriptomic analysis revealed enrichment of ECM degradation and vascular dysfunction pathways, including upregulation of matrix-remodelling and vascular permeability genes such as *Mmp13*, *Mmp3*, and *Ptgs2*, alongside reduced collagen-associated and vascular homeostasis genes. Spatial modelling further showed disrupted collagen–vascular coupling and an association between hypoxia and reduced vessel diameter, consistent with dysfunctional angiogenesis rather than improved perfusion. These changes arose early and independently of tumour size, establishing impaired integrin activation as a central mechanism linking ECM degradation, vascular dysfunction, and sustained hypoxia in aggressive cSCC.

## Introduction

Integrins are central mediators of cell–extracellular matrix (ECM) adhesion that regulate tissue architecture, mechanotransduction, and signal integration within the tumour microenvironment (TME)^1^. Through bidirectional signalling, integrins coordinate cellular responses to biochemical and mechanical cues, thereby influencing tumour initiation, progression, and invasion. Dysregulation of integrin activation has emerged as a key driver of cancer^2^, promoting ECM remodelling, angiogenesis, and adaptation to hypoxic stress across multiple solid tumour types.

Kindlin-1, encoded by *FERMT1*, is an essential co-activator of integrins that facilitates their activation through inside-out signalling^3^. Kindler epidermolysis bullosa (KEB) is a genetic skin disorder caused by loss-of-function mutations in the *FERMT1* gene and is characterised by epithelial fragility and abnormal pigmentation^4,5^ and a strong predisposition to aggressive cutaneous squamous cell carcinoma (cSCC)^6,7^. Beyond its structural role in maintaining epidermal integrity, Kindlin-1 is increasingly recognised as a regulator of keratinocyte signalling and interactions with the microenvironment. Kindlin-1-deficient keratinocytes exhibit enhanced expression of mesenchymal markers, including matrix metalloproteinases (MMPs), and secrete pro-inflammatory cytokines that promote fibrosis and tumour-supportive stromal remodelling^5,8,9^. The ECM is a dynamic and instructive component of the TME, with collagen as its most abundant structural protein and a key determinant of tissue homeostasis^10,11^. In cancer, alterations in collagen organisation, density, and crosslinking contribute to increased tissue stiffness and invasive behaviour^12,13,14^. These changes are often driven by integrin-dependent signalling pathways that regulate both matrix deposition and degradation. For example, collagen crosslinking enzymes of the lysyl oxidase (LOX) family enhance matrix stiffness and have been linked to epithelial-to-mesenchymal transition in advanced cancers^15^. Fibroblasts, as principal producers and remodelers of collagen, further modulate the ECM through secretion of proteolytic enzymes and pro-angiogenic factors^16,17^, reinforcing tumour–stroma crosstalk. Notably, integrin-mediated interactions between tumour cells, fibroblasts, and the ECM are critical for coordinating these processes^1^.

ECM remodelling is tightly coupled to vascular organisation and oxygen availability within the TME^18^. Hypoxia is a hallmark of cSCC progression, with stabilisation of hypoxia-inducible factor-1α (HIF-1α) promoting tumourigenesis and driving angiogenic programmes^19^, including upregulation of vascular endothelial growth factor (VEGF)^20^. In turn, angiogenesis attempts to restore oxygen and nutrient supply^21^ but often results in structurally and functionally abnormal vasculature. In a skin setting a link between angiogenesis and collagen has been reported with expression of collagen-I by fibroblasts and vascular cells having a pro-angiogenic effect in invasive melanoma^22^. In addition, in models of recessive dystrophic EB (RDEB), characterised by absence of collagen VII expression, cSCC presents with increased angiogenesis^23,24^. Emerging evidence suggests that integrin signalling plays a central role in this interplay by regulating endothelial cell behaviour, matrix architecture, and tissue perfusion^2,25^. In skin pathologies and cancers, collagen deposition and organisation have been directly linked to angiogenic responses^22^, while disruptions in ECM composition, such as loss of collagen VII in DEB, are associated with enhanced vascularisation and aggressive tumour phenotypes^23, 24^.

Despite these advances, how defects in integrin activation reshape the spatial and functional coupling between ECM architecture, vascular organisation, and hypoxia within the TME remains poorly understood. In particular, whether impaired integrin signalling can drive coordinated changes in collagen remodelling, angiogenesis, and oxygen distribution to promote tumour progression has not been fully elucidated.

In our previous work^26^, we demonstrated that Kindlin-1 deficiency promotes SCC tumour growth associated with a hypoxic microenvironment, whileenhancing tumour cell invasion through upregulation of MMPs, particularly MMP13. Building on these findings, we hypothesise that loss of Kindlin-1-dependent integrin activation disrupts the coordinated regulation of ECM structure, vascular organisation, and oxygen homeostasis within the TME. Here, using cell culture, three-dimensional tumour spheroids, and in vivo mouse models, we investigate how these processes are integrated in the pro-tumorigenic behaviour of Kindlin-1-deficient cSCC (Fig. 1a).

**Figure 1.**
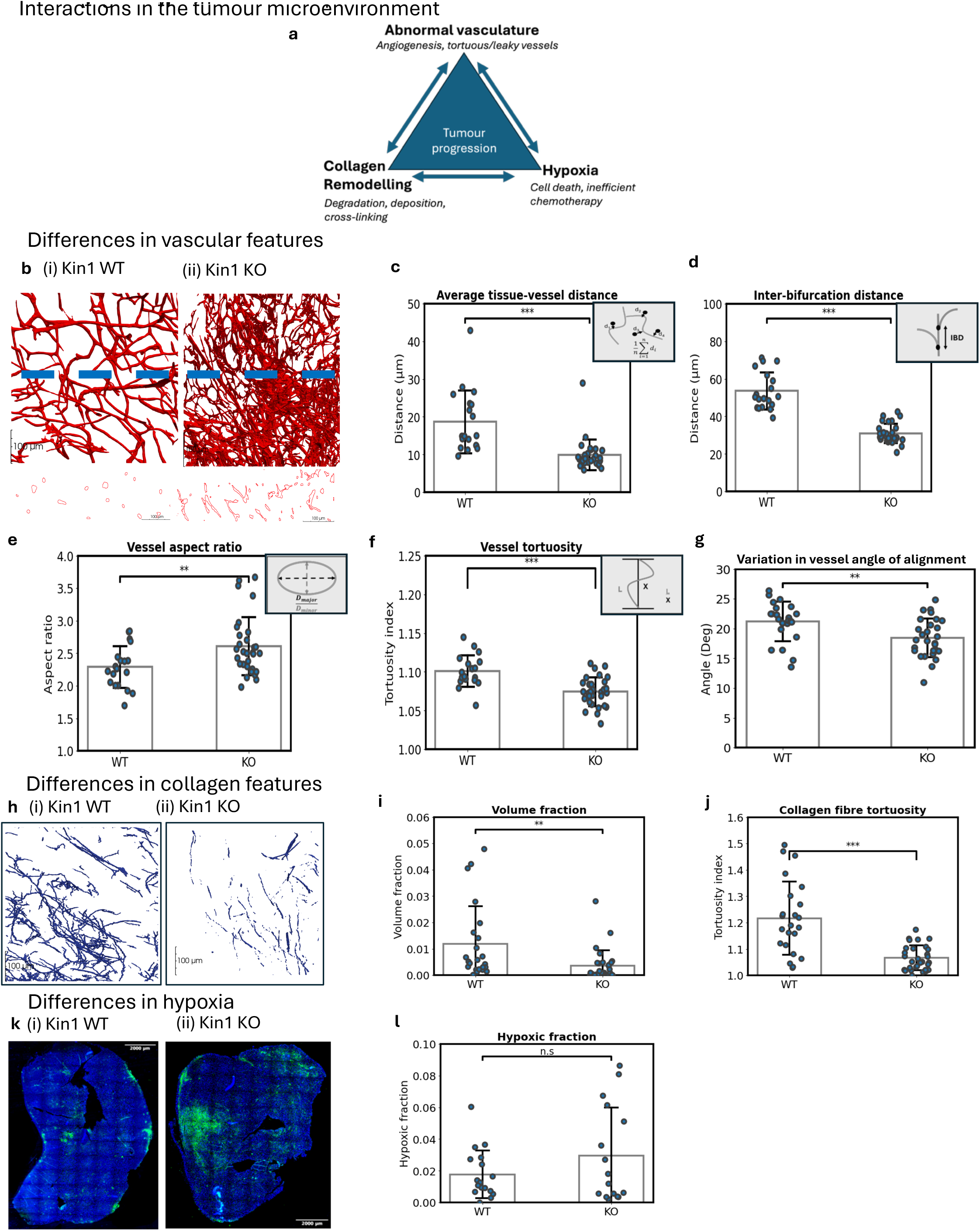
Di4erences in tumour microenvironment phenotypes between Kin1 WT and Kin1 KO tumours. **(a)** Schematic illustration of the principal constituents of the tumour microenvironment (collagen ECM, vasculature, and hypoxia) and their interactions. **(b)** Representative 3D renderings of tumour vasculature from mature (i) Kin1 WT and (ii) Kin1 KO tumours, with corresponding 2D cross-sections shown below (blue dashed line indicates section plane). **(c–g)** Quantitative vessel features derived from 3D image stacks showing significant diMerences between Kin1 WT and Kin1 KO genotypes: **(c)** Mean distance from any point in tissue to the nearest vessel (smaller distance = denser vasculature). **(d)** Mean inter-bifurcation distance. **(e)** Vessel aspect ratio (largest / smallest diameter). **(f)** Vessel tortuosity (path length / Euclidean distance between endpoints). **(g)** Variation in vessel orientation, calculated as the mean angular deviation in x, y, and z. **(h)** Representative 3D collagen I reconstructions from (i) WT and (ii) KO tumours. **(i–j)** Quantitative collagen metrics: **(i)** Collagen volume fraction; **(j)** Collagen-fibre tortuosity. **(k)** Representative immunofluorescence of DAPI (blue) and pimonidazole (green) in (i) WT and (ii) KO tumours. **(l)** Average fraction of pimonidazole-positive (hypoxic) area per 500 µm × 500 µm tile. Data points represent single tiles with stack depths of 155 µm for vasculature, 40 µm for collagen, 1 µm for hypoxia). Stacks were obtained from five WT tumours (n = 3 mice) and four KO tumours (n = 2 mice). Error bars denote ± 1 SD of the mean.

## Results

### Kindlin-1 loss disrupts vascular architecture and extracellular matrix organisation in vivo

Quantitative analysis of 3D vessel networks in cSCC tumours revealed clear structural and organisational diZerences between Kin1 WT and Kin1 KO tumours (Fig. 1b-g). Kin1 KO tumours exhibited a significantly reduced mean tissue-to-vessel distance (Fig. 1c), consistent with a denser microvascular network. Inter-bifurcation distance was also shorter in Kin1 KO tumours (Fig. 1d), a metric previously associated with localised hypoxia^34^, indicating a more compact, highly branching vascular tree.

Vessel morphology diZered markedly by genotype. Kin1 KO tumours showed higher vessel aspect ratios (Fig. 1e), reflecting more anisotropic cross-sections, yet these vessels were paradoxically less tortuous (Fig. 1f) and displayed greater alignment (Fig. 1g). Together, these features point to structurally ordered but functionally abnormal vascular architecture in Kindlin-1-deficient tumours.

ECM composition was similarly altered (Fig. 1h-j). Kin1 KO tumours contained substantially less collagen than Kin1 WT tumours, consistent with reports of Kindlin-1-dependent matrix deposition^26^. Collagen fibres were also significantly straighter in Kin1 KO tumours (Fig. 1j), suggesting that Kindlin-1 loss aZects both collagen quantity and microstructural organisation.

Pimonidazole staining showed a trend towards increased hypoxia in Kin1 KO tumours (Fig. 1k-l), consistent with prior observations^26^.

### Hypoxia-driven transcriptional changes in 2D cultures and spheroids reflect Kindlin-1-dependent responses

Hypoxia has a well-established role aZecting ECM remodelling and abnormal vasculature^26,35,36^. To dissect the mechanisms contributing to these TME alterations, we assessed gene expression in 2D cultures exposed to 3% O₂ for 72 hours. Hypoxic conditions significantly increased expression of *Vegfa* and *Angpt1* (Angiopoietin-1) (Fig. 2a-b), two central regulators of angiogenesis and vascular remodelling^37,38^ in both Kin1 KO and Kin1 WT cells. Kin1 KO cells expressed significantly more *Vegfa* than Kin1 WT cells, while *Angpt1* induction was greater in Kin1 WT cells, suggesting genotype-specific angiogenic programmes in 2D. In contrast, expression of *Col1a1* which encodes collagen 1 (Fig.2c) was significantly decreased under hypoxia in both Kin1 KO and Kin1 WT cells. Induced hypoxia in 2D cultures was confirmed by expression of the hypoxic responsive *Idha* gene (Supplementary Fig. 1).

**Figure 2.**
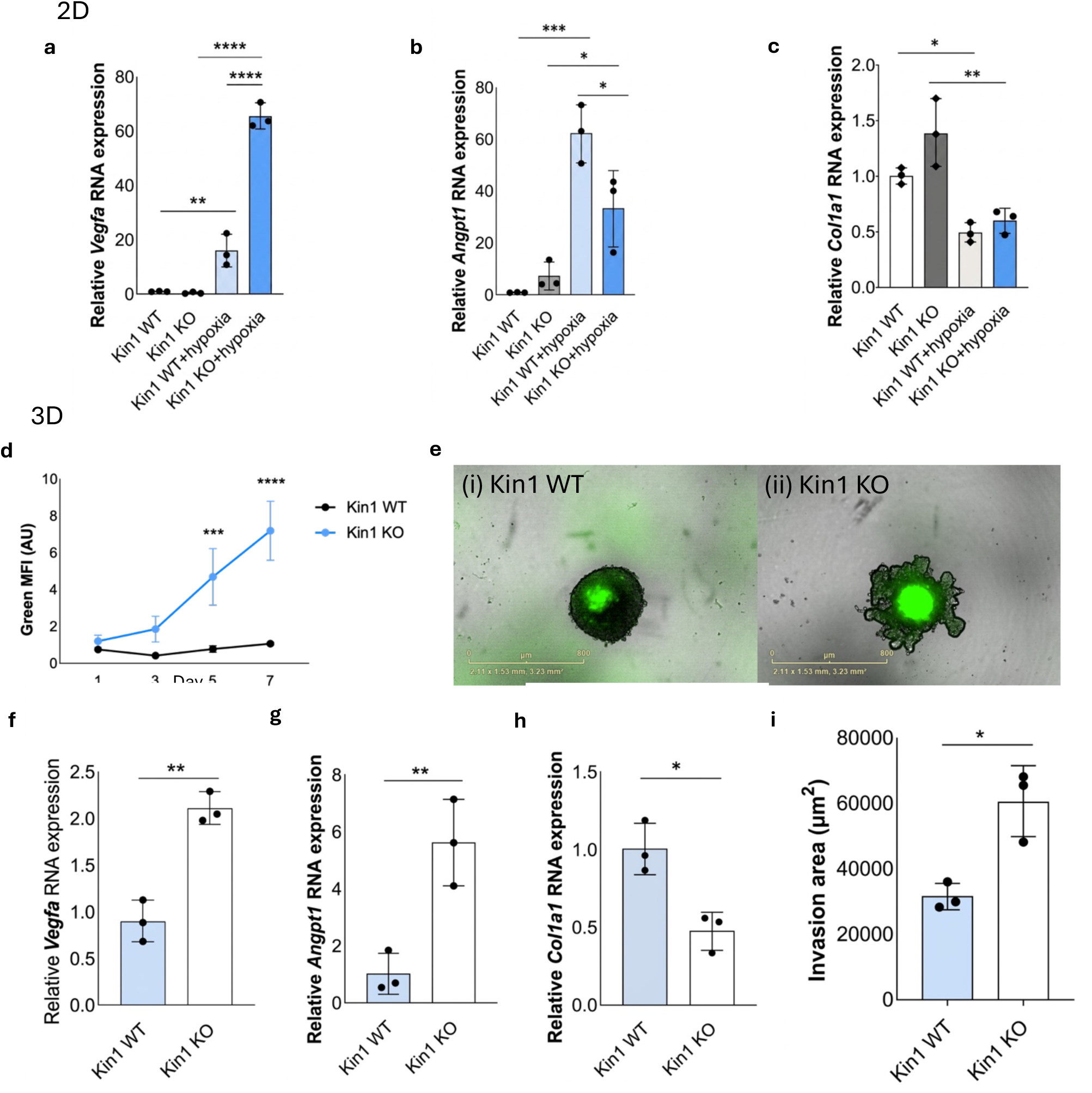
Increased tumorigenic factors in Kin1 KO cSCC cells compared to Kin1 WT. mRNA expression of angiogenic markers (a) *Vegfa* and (b) *Angpt1* and (c) *Col1a1* in normal and hypoxic conditions. (d) Quantification of GFP expression in HRE-GFP-transfected cSCC cells and (e) representative images, scale bar = 800 μm. mRNA expression of angiogenic markers (f) *Vegfa*, (g) *Angpt1*and (h) *Col1a1*in cSCC cells grown as spheroids. (i) Quantification of cSCC spheroid invasion after 7 days. Data obtained from three independent experiments (mean ± SD). p-values were obtained from one-way ANOVA test followed by Tukey post-hoc test; ****p < 0.0001, ***p< 0.001, **p < 0.01 and *p < 0.05.

To investigate if changes in collagen expression were linked to hypoxia-induced remodelling, we used cSCC cells transfected with a hypoxia-responsive element (HRE) expressing GFP^39^. Confirming our previous findings^26^, loss of Kindlin-1 in 3D spheroids showed increased hypoxia by HRE-GFP expression (Fig. 2d). Moreover, consistent with the 2D gene expression findings, vasculature markers were significantly increased in Kindlin-1 deficient spheroids and collagen expression was significantly decreased (Fig. 2f-h). To assess hypoxia and collagen degradation simultaneously, cSCC spheroids were grown in collagen. As shown in Fig. 2i, at the timepoint with the highest HRE-GFP expression (Fig. 2d, 7 days), Kindlin-1 deficient cSCC cells had increased invasiveness which we have previously reported is associated with collagen degradation^26^.

### Kindlin-1 loss drives transcriptomic signatures of ECM remodelling and vascular dysfunction

Previous RNA-seq analysis and in vivo studies showed that Kindlin-1 loss in cSCC leads to the promotion of a highly hypoxic tumour environment, upregulation of epithelial-to-mesenchymal (EMT) markers and a decrease in collagen deposition. We also previously demonstrated that increased invasion of Kindlin-1-deficient cSCC is promoted by upregulation of collagenase-3/matrix metalloproteinase 13 (MMP13)^40^, a key mediator of ECM protein degradation and tumor progression of cSCC^23^.

To determine whether transcriptional changes associated with Kindlin-1 loss reflected the structural alterations observed in the TME, we performed STRING analysis of the top 50 differentially expressed genes between Kin 1 WT and Kin1 KO tumours. The top 50 upregulated and downregulated genes are listed in Supplementary Table 2. This revealed enrichment of biological processes associated with ECM remodelling and vascular regulation in Kin1 KO tumours (Fig. 3a–d).

**Figure 3.**
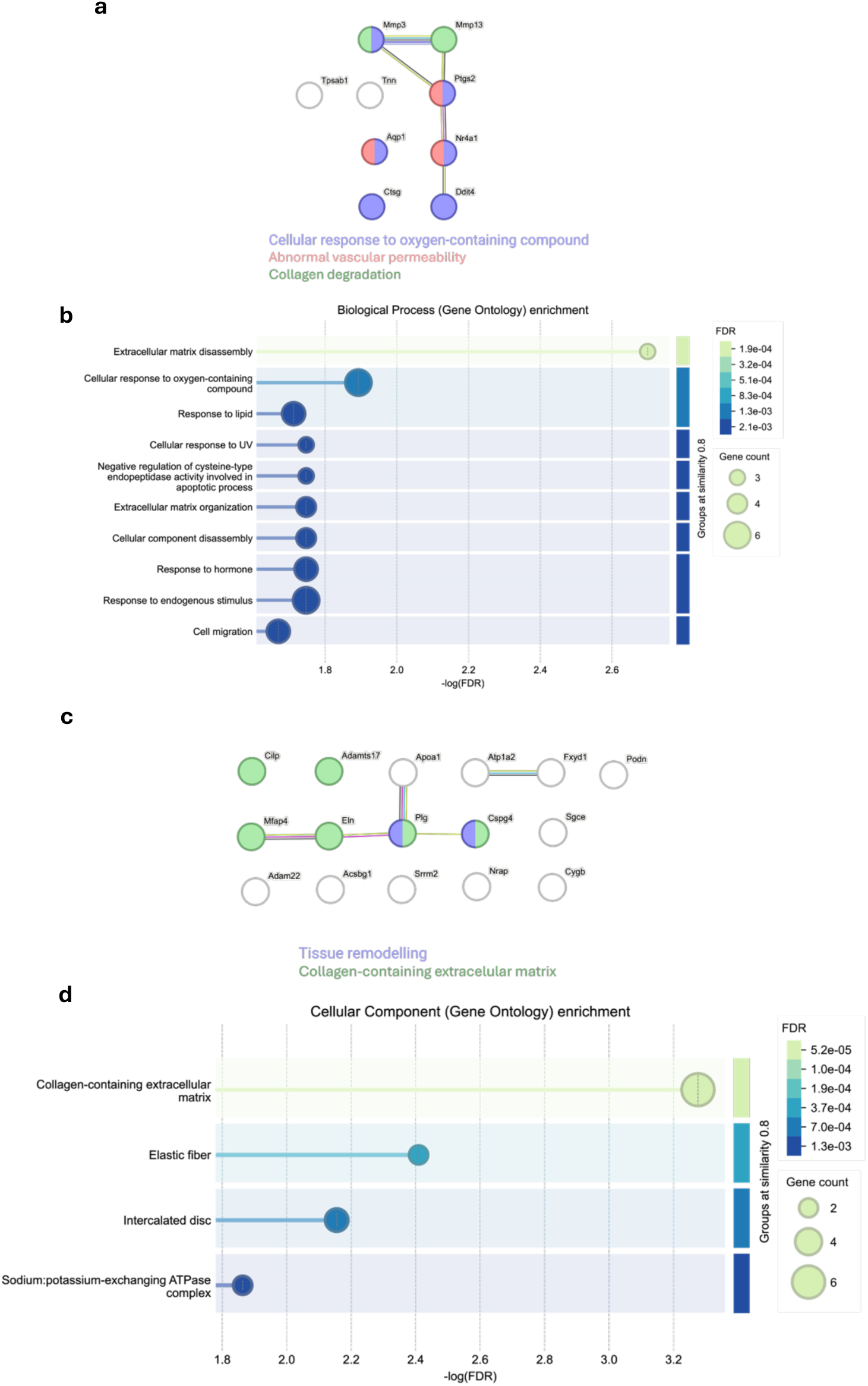
Differential gene expression analysis identifies ECM remodelling and vascular-associated pathways in Kin1 KO tumours. STRING analysis and gene ontology enrichment of selected genes linked to extracellular matrix and vascular features (n=3 tumours). STRING protein–protein interaction and Gene Ontology (GO) enrichment analyses were performed on the top 50 upregulated and downregulated genes identified in Kin1 KO tumours relative to controls (n = 3 tumours per group). (a) STRING interaction network of upregulated genes showing enriched functional clusters associated with cellular responses to oxygen-containing compounds, vascular permeability, and collagen degradation. (b) GO enrichment analysis of upregulated genes demonstrating significant enrichment of pathways related to extracellular matrix disassembly, cellular responses to oxygen-containing compounds, and cell migration. Bubble size indicates gene count and colour represents false discovery rate (FDR). (c) STRING interaction network of downregulated genes identifying clusters associated with extracellular matrix organisation and tissue remodelling. (d) GO enrichment analysis of downregulated genes demonstrating enrichment of extracellular matrix-related components, including collagen-containing extracellular matrix and elastic fibre organisation. Bubble size indicates gene count and colour represents FDR.

Among the upregulated genes, several were associated with proteolytic matrix degradation and vascular permeability, including *Ctsg*, *Tpsab1*, and the matrix metalloproteinases *Mmp13* and *Mmp3*, consistent with enhanced ECM breakdown and tissue remodelling. In contrast, downregulated genes included multiple factors linked to collagen-containing ECM and vascular homeostasis, including *Smoc1*, *Cilp*, *Adamts17*, *Eln*, and *Mfap4*, as well as genes associated with actin organisation and cardiovascular physiology.

Transcriptional signatures associated with vascular dysfunction and increased permeability, including *Ptgs2*, *Aqp1*, *Nr4a1*, and *Plac8*, were also identified, consistent with a model in which Kindlin-1 loss promotes both ECM destabilisation and abnormal vascular function. Together, these findings indicate coordinated disruption of matrix organisation and vascular integrity, contributing to the highly pro-tumorigenic microenvironment observed in Kin1 KO tumours. A broader representation of the top 50 differentially expressed genes is provided in Supplementary Fig. 2.

### Kindlin-1 loss uncouples the relationships between hypoxia, vasculature, and collagen *in vivo*

Given these hypoxia-associated transcriptional changes, we next investigated how hypoxia spatially relates to vascular and collagen organisation *in vivo*. Mixed-effects modelling demonstrated that in Kin1 KO tumours, hypoxia was strongly associated with reduced vascular area in 2D cross-sections (β = −0.116; *p* < 0.001; Fig. 4a). A similar, although not significant, trend was shown between hypoxia and vessel volume fraction in 3D reconstructions (β = −0.353; *p* = 0.152; Supplementary Fig. 3a). Hypoxia was also associated with reduced vessel diameter in 3D reconstructions (β = −0.594; *p* = 0.037; Fig. 4b), indicating localised vessel thinning. In addition, hypoxia showed a non-significant trend towards decreased inter-bifurcation λ, a measure of vessel length-to-diameter ratio (β = −0.312; *p* = 0.057; Fig 4c). Smaller λ values are associated with abnormal vessel morphology and have been proposed to bias haematocrit distribution^41^. Importantly, none of these associations between vasculature and hypoxia were observed in Kin1 WT tumours.

**Figure 4.**
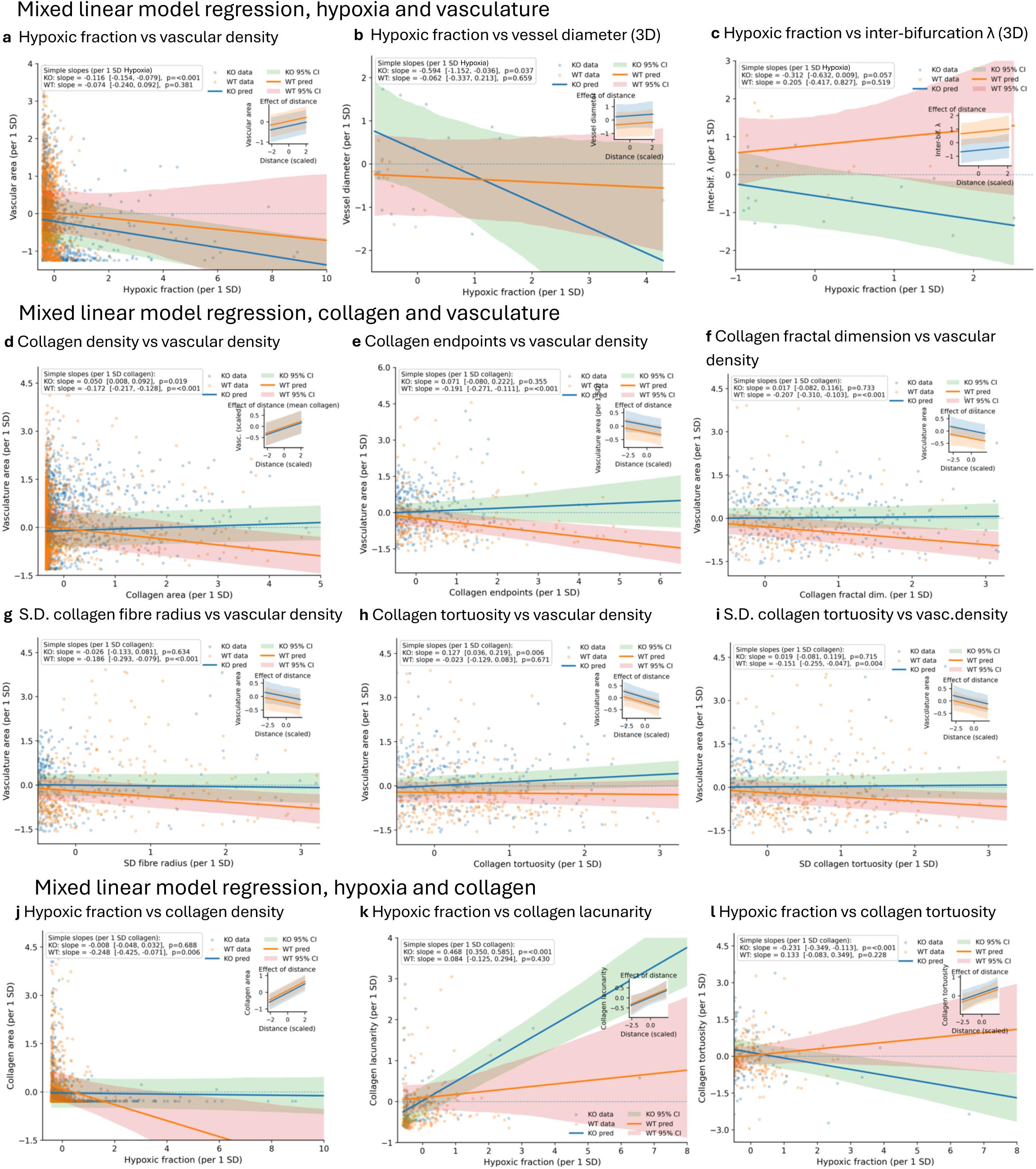
Relationships between vasculature, collagen, and hypoxia in Kin1 WT and Kin1 KO tumours. Each scatter plot shows the relationship between two microenvironmental features across individual image tiles or 3D volumes; solid lines represent genotype-specific regressions (KO, blue; WT, orange) with shaded 95% confidence intervals. Insets show the eMect of radial distance, confirming independence from spatial gradients. **(a–c)** Vascular features vs hypoxia: (a) vascular area, (b) vascular volume, and (c) mean vessel diameter; KO tumours show negative associations, whereas WT shows no significant associations. **(d–i)** Vascular-collagen relationships across collagen (d) density (KO positive, WT negative), (e) endpoints (WT negative, KO no significant association), (f) fractal dimension (WT negative, KO no significant association), (g) fibre-radius SD (WT negative, no significant KO association), (h) tortuosity (KO positive, WT none), and (i) tortuosity SD (WT negative, KO no significant association). **(j–l)** Collagen features vs hypoxia across collagen (j) area (WT negative, KO no significant association), (k) lacunarity (KO positive, WT no significant association), and (l) tortuosity (KO negative, WT no significant association). Data are combined from five tumours from three Kin-1 WT mice and four tumours from two Kin-1 KO mice.

Collagen-vascular relationships were also genotype-specific. In Kin1 WT tumours, increased collagen area (β = −0.172; *p* <0.001), collagen endpoints (β = −0.191; *p* <0.001), fractal dimension (β = −0.207; *p* <0.001), and fibre-radius variability (β = −0.172; *p* <0.001) were each associated with reduced vascular area (Fig. 4d-i). In contrast, these associations were largely absent or reversed in Kin1 KO tumours, indicating disrupted ECM–vascular coupling following Kindlin-1 loss. No associations were observed between vascular inter-bifurcation λ and collagen density in either genotype (Supplementary Fig. 3b).

Hypoxia-collagen relationships showed a similar divergence between genotypes. In Kin1 WT tumours, hypoxia was associated with local collagen loss (Fig. 4j). In contrast, hypoxia in Kin1 KO tumours was associated with increased collagen lacunarity and straighter fibres (Fig. 4k), suggesting that Kindlin-1 loss changes the structural response of collagen to hypoxia.

### Collagen-vascular interactions evolve with tumour maturity and diverge early in Kin1 KO tumours

To investigate how the TME organisation evolves with progression, we compared collagen and vascular features in size-matched small (5 mm) and medium (10 mm) Kin1 WT and Kin1 KO tumours. As Kindlin-1-deficient tumours are known^26^ to grow more rapidly, size-matching enables assessment of whether phenotypic differences arise independently of tumour size.

At early stages (5 mm), Kin1 KO tumours exhibited increased collagen density compared to Kin1 WT tumours, although WT tumours showed greater variability (Fig. 5a). By 10 mm, collagen density decreased in both genotypes, with Kin1 KO tumours becoming less collagen-dense than WT. In contrast, vascular density was comparable between genotypes at 5 mm (Fig. 5b), but increased in Kin1 KO tumours at 10 mm, accompanied by greater variability.

**Figure 5.**
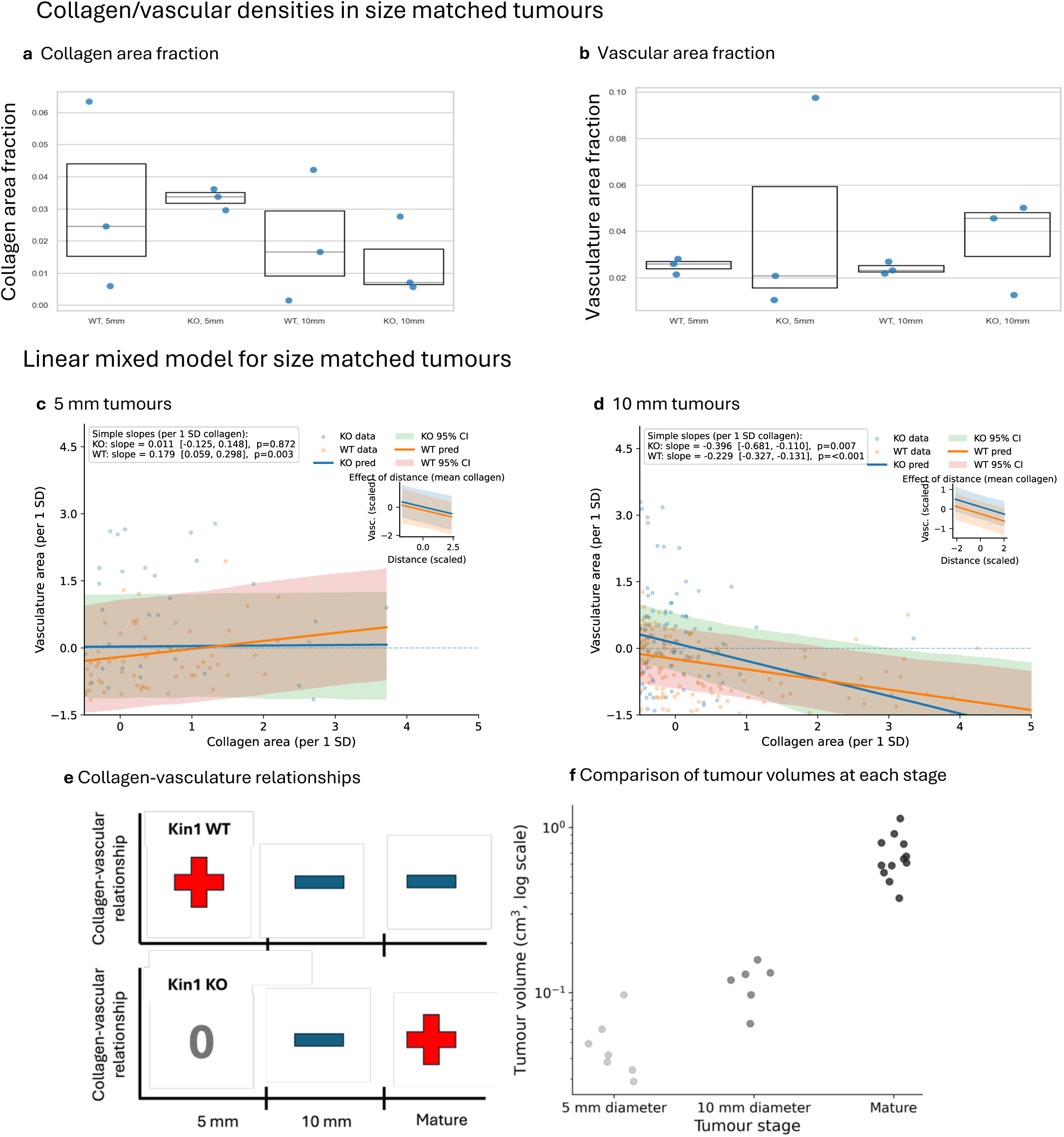
Analysis of collagen and vasculature in size-matched Kin1 WT and Kin1 KO tumours. (a) Average collagen fraction per 2D tumour tile for Kin1 WT and Kin1 KO tumours with average diameters of 5mm 10mm. (b) Average vasculature fraction per 2D tumour tile for Kin1 WT and Kin1 KO tumours with average diameters of 5mm and 10mm. (c-d) Scatter plots shows the relationships between collagen density and vascular density across image tiles for (c) 5mm tumours and (d) 10 mm tumours; solid lines represent genotype-specific regressions (Kin1 KO, blue; Kin1 WT, orange) with shaded 95% confidence intervals. Insets show the eMect of radial distance, confirming independence from spatial gradients. (e) Summary illustration of relationships between collagen and vascular densities in in-vivo WT and KO tumours with 5mm and 10 mm diameters and in mature tumours (Positive, red; Negative, blue, No relation, grey). (f) Log-scale distributions of tumour volumes from 5mm and 10 mm diameters and in mature tumours. Data are combined from two 5mm tumours from two Kin-1 WT mice, four 5mm tumours from two Kin-1 KO mice, two10mm tumours from one Kin-1 WT mouse and three 10 mm tumours from two Kin-1 KO mice.

These structural changes were associated with early divergence in collagen–vascular coupling. In small (5 mm) tumours, Kin1 WT tumours displayed a positive correlation between collagen and vascular density, consistent with coordinated matrix-guided angiogenesis, whereas this relationship was absent in Kin1 KO tumours (Fig. 5c). By 10 mm, both genotypes exhibited negative correlations (Fig. 5d), suggesting a shift towards decoupling as tumours progress.

Integration with our analysis of mature tumours shown in Fig. 2 indicates that this coordinated relationship is maintained in Kin1 WT tumours but breaks down in Kin1 KO tumours (Fig. 5e), consistent with progressive disruption of ECM–vascular organisation. Importantly, comparison of tumour volumes (Fig. 5f) highlights that 5 mm and 10 mm tumours represent relatively early-stage lesions compared with mature tumours.

Despite this size-matching, phenotypic differences between genotypes were evident. Kin1 KO tumours showed disrupted collagen–vascular coupling at 5 mm and increased vascular density at 10 mm, indicating that alterations in TME organisation arise independently of tumour size. Although the sample size limits statistical power at early stages, these findings suggest that Kindlin-1 loss perturbs ECM-guided angiogenesis from the earliest stages of tumour development and promotes aberrant vascular expansion with progression.

## Discussion

Our integrative analysis of 2D cultures, 3D spheroids, and in vivo tumours demonstrates that loss of Kindlin-1 disrupts the organisation and function of the TME in cSCC. Given the essential role of Kindlin-1 in integrin activation^3^, these findings strongly implicate impaired integrin signalling as a central regulator of coordinated alterations in ECM architecture, vascular organisation, and oxygen homeostasis. Kindlin-1 deficiency in cancer cells promotes a hypoxic state, which may both drive and be reinforced by vascular dysfunction, while simultaneously accelerating ECM degradation through elevated MMP-13^26^ and suppressing collagen I transcription. The resulting microenvironment is characterised by a structurally simplified and porous collagen network alongside a structurally ordered but functionally compromised vasculature.

These observations support a model in which loss of integrin activation uncouples the reciprocal interactions between tumour cells and their microenvironment. Rather than acting through independent pathways, ECM remodelling and angiogenesis emerge here as interdependent processes that are normally coordinated through integrin-mediated mechanotransduction. In the absence of Kindlin-1, this coordination breaks down, giving rise to a dual but interdependent mechanism of TME remodelling (Fig. 6): one driven by loss of ECM integrity and mechanical cues, and the other by hypoxia-driven, maladaptive angiogenesis. This framework provides a plausible, mechanistic explanation for the increased invasiveness observed in our models and may explain the aggressive cSCC phenotypes arising in Kindler epidermolysis bullosa (KEB). It is well-established that patients with KEB present defects in integrin activation, affecting signal transduction between the local ECM and cells^34–36^.

**Figure 6.**
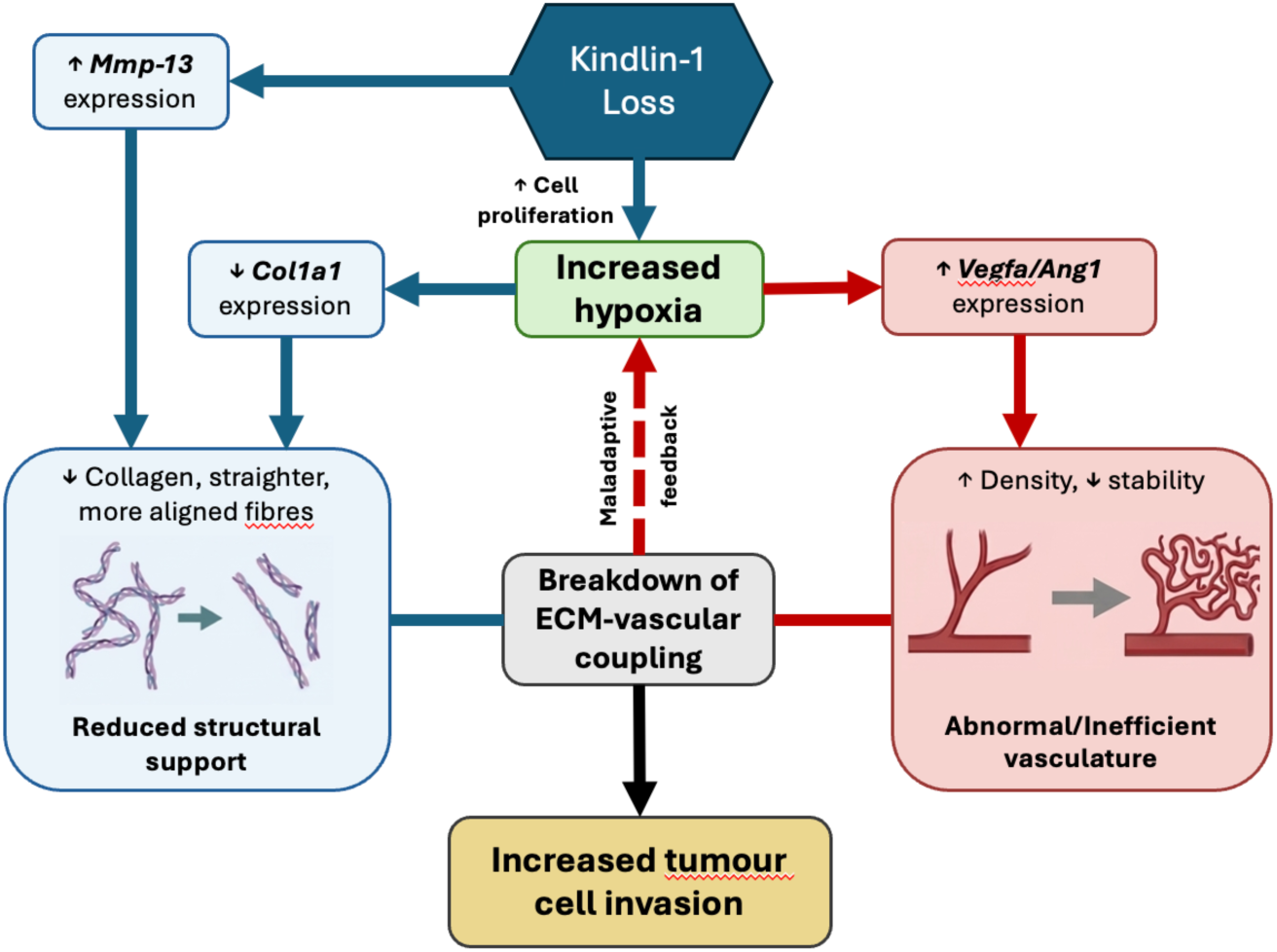
Proposed two-pathway mechanism by which Kindlin-1 knockout remodels the tumour microenvironment. Loss of Kindlin-1 is associated with two pathways aMecting tumour microenvironment organisation. **Left (blue):** Reduced Kindlin-1 function is associated with increased *Mmp-13* expression, promoting collagen degradation. In parallel, elevated cell proliferation elevates oxygen demand, contributing to local hypoxia, which in turn downregulates *Col1a1* expression. Together, these changes alter extracellular-matrix architecture resulting in a structurally simplified matrix with reduced support for vascular organisation. **Right (red):** Increased hypoxia is accompanied by upregulation of *Vegfa* and *Angpt1*, consistent with enhanced but aberrant angiogenic signalling and the formation of structurally abnormal and functionally ineMicient vasculature. The combined eMects of these pathways are associated with disruption of the normal coupling between extracellular matrix organisation and vascular structure observed in wild-type tumours. These observations support a model in which hypoxia and vascular dysfunction reinforce one another, contributing to a persistently hypoxic and structurally disorganised tumour microenvironment that is permissive for tumour cell invasion.

At the molecular level, our transcriptomic analyses further support widespread disruption of integrin-regulated pathways. Upregulated genes in Kindlin-1-deficient tumours are enriched for factors associated with vascular permeability and ECM degradation (e.g. *Ctsg, Tpsab1*), including proteases and matrix-remodelling enzymes, while downregulated genes are enriched for pathways associated with maintaining vascular homeostasis and ECM organisation, suggesting a potential influence of these features in the highly pro-tumourigenic tumour environment observed in Kin1 KO tumours. These changes are consistent with a shift from a mechanically structured to a permissive matrix environment. Together, these data suggest that loss of Kindlin-1 drives net matrix loss through both enhanced degradation and reduced synthesis. Given that integrins regulate both matrix assembly and proteolytic remodelling, these findings suggest that loss of Kindlin-1 skews the balance of ECM turnover towards degradation while impairing the establishment of organised collagen architecture.

Expression data from mouse tumours show that the reduced collagen content in Kin1 KO tumours can be linked to reduced expression of genes related to collagen-containing ECM (e.g. *Smoc1, Cilp, Adamts17, Plg, Eln, Mfap4*). In addition, gene expression changes in tumour spheroids support this interpretation, indicating reduced ECM deposition and increased ECM degradation. Moreover, analysis of collagen in 3D images of mouse cSCC indicates that Kindlin-1 loss results in straighter, more aligned collagen fibres which may facilitate invasion through straighter fibre “tracks”^45^. Consistent with this, Kin1 KO spheroids exhibiting higher hypoxia also showed increased invasive capacity. The increased hypoxic state we have consistently observed in Kin1 KO cSCC cells and tumours supports a role for hypoxia as a key driver for ECM remodelling. Hypoxia is known to induce the expression of matrix metalloproteinases, known ECM degraders reported in the literature^26,46,47^, increasing the invasive capabilities of cancer cells in order to migrate.

Our data also identify hypoxia as a central component of this rewiring of the TME. Kindlin-1-deficient tumours exhibit sustained hypoxia despite increased vascular density, indicating a decoupling between angiogenesis and functional perfusion. This phenotype is consistent with the formation of immature and inefficient vasculature, as reflected by increased expression of pro-angiogenic factors such as *Vegfa* and *Angpt1*. Furthermore, transcriptomic signatures of increased vascular leakiness and abnormal vascular function (e.g. *Ptgs2, Aqp1, Nr4a1, Plac8*) further support a model in which Kindlin-1 loss destabilises the vasculature at both structural and functional levels.

Mechanistically, disruption of Kindlin-1-dependent integrin activation may impair downstream mechanotransduction pathways involved in matrix adhesion, endothelial behaviour, and vessel stabilisation such as FAK/Src signalling^48^. In parallel, sustained hypoxia may promote compensatory HIF-dependent angiogenic signalling, although the resulting vasculature appears functionally inadequate. Together with increased ECM degradation and altered collagen organisation, these changes are likely to impair tissue perfusion and oxygen delivery. The persistence of hypoxia in this context therefore supports a model in which hypoxia and vascular dysfunction reinforce one another, contributing to a progressively disorganised and metabolically stressed TME.

Consistent with this, despite increased vascular density, Kin1 KO tumours remain hypoxic, in agreement with previous studies showing a lack of correlation between hypoxia and the density of the microvasculature in head and neck SCC^49^. Feature analysis of mature tumours shows that hypoxic regions contain thinner vessels and reduced vascular area. These observations provide spatial evidence of vascular dysfunction, indicating that newly formed vessels are structurally inefficient and fail to relieve oxygen deprivation. Such a phenotype is consistent with pathological angiogenesis observed in many hypoxic, treatment-resistant solid tumours^50^.

The altered ECM-vascular relationships we observe further underscore the role of Kindlin-1 in coordinating mechanical and metabolic features of the TME. Whereas Kin1 WT tumours show evidence of collagen-guided sprouting^51^ and later collagen-mediated vascular restriction, Kin1 KO tumours lose these regulatory cues early, with consequences for vessel patterning and tumour invasion. This aligns with prior work showing collagen stiffness and alignment shape tumour progression^52^ and that endothelial sprouting typically follows collagen fibres^53^.

Although this study focuses on genetic loss of Kindlin-1, the resulting phenotype support a more general role for integrin activation in maintaining TME homeostasis. By coordinating ECM organisation, vascular function, and oxygen distribution, integrin signalling can act as a central integrator of mechanical and metabolic cues within tumours. Its disruption leads to a collapse of this coordinated system, producing a microenvironment that is structurally permissive, metabolically stressed, and highly pro-invasive. These insights may extend beyond KEB-associated cSCC to other solid tumours in which integrin signalling and ECM remodelling are dysregulated.

Future work should directly assess how impaired integrin activation affects vascular perfusion and oxygen delivery dynamics, and whether restoring integrin signalling can normalise ECM architecture and vascular function. Extending these analyses to patient-derived samples will be essential to determine the clinical relevance of these mechanisms and to evaluate whether targeting integrin-dependent pathways may provide therapeutic benefit through simultaneous modulation of tumour mechanics, angiogenesis, and hypoxia.

## Methods

### In vitro studies

GFP-labelled Kindlin-1 wild type (Kin1 WT) or Kindlin-1 knockout (Kin1 KO) SCC cell lines were generated and grown as previously described^26,27, 28^. Cells were routinely tested for mycoplasma every month with a Mycoalert® Mycoplasma detection kit (Lonza, United Kingdom). For assessment of hypoxia expression, stable cell lines were generated in Kin1 WT and Kin1 KO SCC cells using a plasmid containing 5 hypoxia-responsive element (HRE) sequences in tandem to a CMV promoter (https://www.addgene.org/46926/), kindly provided by Friedmann Kiefer (European Institute for Molecular Imaging, Münster)^29^. Transfections were performed using jetOPTIMUS® transfection reagent (Polyplus, France), following manufacturer’s instructions (1 mg of DNA). After transfection, cells were selected by incubation with 1.5 mg/ml geneticin (Thermo Fisher Scientific, Waltham, MA, USA) for 24 hours. For hypoxia studies, SCC cells (0.5x10^6^) were plated in 6-well plates and after 24 hours, cells were incubated for 72 hours in a humidified incubator at 37°C in 3% v/v oxygen prior to RNA extraction. For spheroid studies, SCC cell lines were plated as specified in our previous studies^26^ and grown for 7 days.

### In vivo studies

All experiments were carried out in compliance with UK Home Office regulations (PP7510272) and were approved by the University of Edinburgh Animal Welfare and Ethical Review Body (PL05-21). To mark the vasculature Cdh5-CreER^T2^; Rosa26^lsl-tdT^ mice were treated with tamoxifen (100 mg/kg) by oral gavage daily for 5 days to induce expression of TdTomato VE-cadherin (Cdh5)^30^. GFP-labelled Kin1 WT or Kin1 KO SCC cells were injected subcutaneously bilaterally (0.25x10^6^ in a volume of 100 μl) and tumours were allowed to establish until they reached a suitable size (∼15 mm diameter or 5 mm/10 mm if specified). 1 hour prior to cull, mice were injected intravenously with Hypoxyprobe™-1 pimonidazole HCl (60 mg/kg; Hypoxyprobe, Burlington, MA, USA) and perfuse fixed. Tumours were collected into 4% paraformaldehyde and after 24 hours, tissue was stored in PBS, embedded in plastic molds with 2% low-melting point agarose and 500 μm sections cut using a Vibratome Leica VT1200 S (Leica Biosystems, Germany).

### Image acquisition

Two-photon fluorescence images were acquired using a custom-built multi-modal microscope setup. A picoEmerald S (APE, Berlin, Germany) laser provided both a tunable pump laser (set to 797.4 nm, 2 ps, 80 MHz repetition rate) and a spatially and temporally overlapped Stokes laser (1031 nm, 2 ps, 80 MHz repetition rate). The output beams were inserted into the scanning unit of an Olympus FV1000MPE microscope using an Olympus XLPL25XWMP N.A. 1.05 objective. Backscattered two-photon fluorescence were collected using a short-pass 690 nm dichroic mirror and IR cut filter (Olympus). A series of filters and dichroic mirrors were then used to deconvolve the different emission signals onto two photomultiplier tubes (PMT). Emissions were filtered using: 570dcxr, FF440/520-Di01 (Semrock) and FF01-400/40 (Semrock) for SHG; 570dcxr, FF440/520-Di01 (Semrock) and FF01-510/84 (Semrock) for GFP and tdTomato signals using 570dcxr, and T640lpxr and HQ610/75m. All filters Chroma unless otherwise stated.

Laser powers after the objective were measured up to 40 mw for the pump and Stokes laser. Images were recorded by FV10-ASW software (Olympus) using 1024 × 1024 pixels over a 509 × 509 μm field of view, with a pixel dwell time of 4 μs. Tumour slices were imaged using the multi-area time-lapse function, whilst image stacks of 155 slices were recorded with a z interval of 1 μm, PMT voltages were adjusted with depth to maintain brightness.

Further imaging was performed using a Zeiss Axio Scan Z1 slide scanning microscope (Carl Zeiss UK, Cambridge, UK) with Fluar or Plan Apochromat objective lenses (2.5x, 5x, 10x, 20x, 40x), a Zeiss Colibri 7 LED light source and a Zeiss Axiocam 506m monochrome CCD camera. Fluorescence filters comprise of Zeiss 90 HE, 92 HE, 96 HE, 38 HE and 43 HE sets. Image capture was performed using Zen Blue Slidescan acquisition software (version 3.4.91).

### Tissue processing

Five sections from the 500 μm brain slices (4 μm thickness every 10 μm) were deparaffinised and rehydrated with xylene and decreasing gradients of ethanol. For detection of hypoxic areas, antigen retrieval was performed with 0.1 M citrate buffer (pH 6.0) 100°C and non-specific binding was blocked with serum-free protein solution for 10 minutes at room temperature. Then, samples were stained with FITC-conjugated mouse monoclonal pimonidazole antibody (1:100; Hypoxyprobe, Burlington, MA, USA) and goat polyclonal mCherry antibody (1:500; Amsbio, United Kingdom) overnight at 4°C. Samples were washed with TBS-T, incubated with donkey secondary anti-goat AF594 (1:500; Thermo Fisher Scientific, Waltham, MA, USA) and mounted with VECTASHIELD® mounting medium with DAPI (Vector Labs, Newark, CA, USA). Image acquisition was performed with Zeiss Axioscan Z1 to acquire fluorescent images with DAPI, FITC and AF594 channels (10×). Background signal from all channels was removed by thresholding samples without antibody incubation on Zen blue software (Version 3.4.91.00000).

### RNA extraction and RTqPCR

SCC tumours (30 mg) were thoroughly homogenised by sterile scalpels and lysed with 0.5 ml TRIzol™ (Thermo Fisher Scientific, Waltham, MA, USA). Samples were incubated for 10 minutes on a tuberoller at room temperature. Chloroform (0.2 ml per 1 ml) was added, the sample was shaken vigorously and incubated for 3 minutes at room temperature. Samples were centrifuged for 5 minutes at 12,000 x g at 4°C. The aqueous phase (colourless upper phase) was transferred to a fresh tube for extraction. For extraction, total RNA from cell pellets was isolated using a RNeasy® Mini Kit (QIAGEN, Venlo, Netherlands) according to the manufacturer’s instructions, as previously described^26^. qPCR was performed based on SYBR™ Select Master Mix protocol (Thermo Fisher Scientific) using the StepOne Real-Time PCR System (Applied Biosystems, Waltham, MA, USA) and analysed as previously described^26^. Primer sequences obtained from IDT (Coralville, IA, USA) are listed in Supplementary Table 1.

### STRING analysis

RNA-Sequencing (RNA-Seq) was previously performed on Kin1 WT and Kin1 KO tumour samples (*n*= 3 tumours)^27^. From this, the top 50 upregulated and downregulated genes were selected and used as input on STRING (https://string-db.org/) for a multiple protein analysis. Features related to the terms ‘collagen, tissue remodelling and vasculature’ were selected for visualization. To narrow genes related to ECM, genes related to the terms ‘extracellular matrix, protease, integrins and vascular response’ were selected for visualization. Cellular Component (Gene Ontology) enrichment graphs were obtained from the protein network of the genes used as input.

### Image processing

A cross-sectional image of vasculature from each mouse tumour was subdivided into 500 μm × 500 μm tiles for two-dimensional (2D) analysis. For three-dimensional (3D) analysis of vascular architecture, multiple stacks of 509 × 509 μm tiles were acquired for each tumour with 155 images recorded with a depth 1 μm between images.

Cross-sectional SHG collagen images from the same tumour tissue were divided into tiles for 2D analysis, and multiple stacks of tiles were acquired for 3D analysis. Collagen and vasculature tiles were extracted from corresponding spatial regions of the tumour cross-section. As SHG signal intensity attenuates with imaging depth, only the upper 40 μm of collagen images from each stack were retained for analysis.

The same tumours were stained with pimonidazole to identify hypoxic regions. Two-dimensional cross-sectional images were acquired and spatially registered to the corresponding vasculature and collagen images using the *Big Warp* plugin for ImageJ.

Segmentation of vasculature, collagen, and hypoxic regions was performed using the Labkit plugin for ImageJ^31^. Segmented vasculature and collagen regions from 3D image stacks were used to define vessel and collagen surfaces using VTK software. Centrelines were extracted from these surfaces and represented as graph structures using *Kimimaro* software (https://github.com/seung-lab/kimimaro).

For mature tumours, vasculature, collagen, and hypoxia data were extracted from, a total of five tumours derived from three Kin-1 WT mice and four tumours derived from two Kin-1 KO mice.

For the tumour diameter-matched study, data was extracted from two 5mm diameter tumours derived from two Kin-1 WT mice, four 5mm tumours from two Kin-1 KO mice, two10mm tumours from one Kin-1 WT mouse and three 10 mm tumours from two Kin-1 KO mice.

### Feature metrics

From 3D vasculature and collagen image stacks, we calculated average vessel diameter, average collagen fibre diameter, vessel aspect ratio, tissue–vessel distance, tissue–fibre distance, inter-bifurcation distance, and length-based tortuosity for both vessels and collagen fibres. Full definitions and formulations of these metrics are provided in Chapter 3 of Enjalbert (2023)^32^.

For each collagen fibre, orientation was estimated in MATLAB as Euler angles (𝜃*_i,x_*, 𝜃*_i,y_*, 𝜃*_i,z_*) describing the longest principal axis of the fitted ellipsoid about the x-, y-, and z-axes. Angles were mapped to a 0–180^∘^ axial representation and transformed using a double-angle mapping,

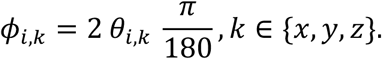

For each component, the mean resultant length was computed as

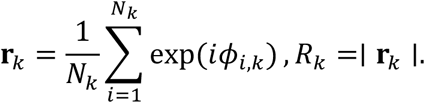

The angular deviation for component 𝑘 is then defined as:

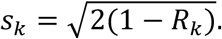

Finally, overall orientation variability is quantified as:

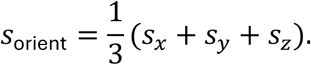

Two-dimensional collagen metrics, including area fraction, total fibre length, lacunarity, number of endpoints, number of branch points, hyphal growth unit (HGU), fibre curvature, and box-counting fractal dimension, were quantified using ImageJ in combination with The Workflow Of Matrix BioLogy (TWOMBLI) ^33^. Fibre alignment was assessed using the coherency metric calculated with the OrientationJ plugin for ImageJ as part of the TWOMBLI workflow. Two-dimensional fibre width, defined as the mean distance from all points on a fibre skeleton to the nearest fibre boundary pixel, and 2D length-based tortuosity were quantified using MATLAB.

### Statistics

Statistical diZerences were assessed using Welch’s two-sample *t*-test, which does not assume equal variances. Tests were two-tailed unless otherwise stated.

TME feature relationships were analysed using linear mixed-eZects models in Python, with one feature (𝑌*_ij_*) modelled as a function of another (𝑋*_ij_*). A covariate term (𝐶*_ij_*) was included to assess the eZects of either genotype (Kin1 WT vs Kin1 KO) or tumour size (5 mm vs 10 mm diameter), together with an interaction term (𝑋*_ij_*𝐶*_ij_*). Radial distance from the tumour centre (𝐷*_ij_*) was included as an additional fixed eZect. Tumour identity was included as a random intercept 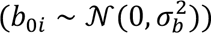.

For observation 𝑗 in tumour 𝑖, the model can be written as:

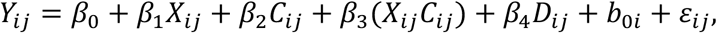

Where residual error 𝜀*_ij_* ∼ 𝒩(0, 𝜎.). Models were fitted using maximum likelihood estimation.

## Acknowledgements

We would like to thank Morwenna Muir and Fraser Laing for their involvement in animal experiments. We would like to thank the service provider Histology Research Service at the Institute of Genetics & Cancer, University of Edinburgh for the processing of mouse tissue for IHC. This work was supported by grants from DEBRA International (Brunton 2), the Cancer Research UK Scotland Centre Edinburgh (CTRQQR-2021\100006) and UKRI Frontier Research grant number EP/X025705/1. MOB acknowledges funding from the European Union’s Horizon 2020 research and innovation program under Grant 801423.

## Author CRediT Statement

Conceptualization: DH, GC, VGB, MOB

Investigation: DH, GC, ML, MF, RE

Formal analysis: DH, GC, RE

Funding acquisition: VGB, MOB

Writing-original draft: DH, GC

Writing-review and editing: DH, GC, ML, VGB, MOB

