## Supplementary Tables and Figures for "Kindlin-1 loss disrupts vascular and extracellular matrix organisation to sustain hypoxia in cutaneous squamous cell carcinoma"

### Supplementary materials

#### Tables

|  | Forward | Reverse |
| --- | --- | --- |
| *Vegf-a* | AAAAACGAAAGCGCAAGAAA | TTTCTCCGCTCTGAACAAGG |
| *Col1a2* | TGCAGTAACTTCGTGCCTAG | ACGTGGTCCTCTGTCTCCA |
| *Ang-1* | GGGGGAGGTTGGACAGTAA | CATCAGCTCAATCCTCAGC |
| *Ldha* | TGTGGCAGACTTGGCTGAGA | CTGAGGAAGACATCCTCATTGATTC |
| *Actb* | GGCTGTATTCCCCTCCATCG | CCAGTTGGTAACAATGCCATGT |

**Supplementary Table 1: Mouse primers used for RT-qPCR.**

|  | Upregulated | Downregulated |
| --- | --- | --- |
| 1 | *F630028O10Rik* | *Neu2* |
| 2 | *Tpsab1* | *Ttn* |
| 3 | *Apcdd1* | *Klra17* |
| 4 | *Cpa3* | *Smoc1* |
| 5 | *Rcan1* | *Adamts17* |
| 6 | *Rgs18* | *Tnrc6b* |
| 7 | *Islr* | *Atp1a2* |
| 8 | *Aldh3a1* | *Tnnt3* |
| 9 | *Plac8* | *Myh4* |
| 10 | *Aqp1* | *Fxyd1* |
| 11 | *Gch1* | *Dusp13* |
| 12 | *S100a9* | *Cdnf* |
| 13 | *Ehf* | *Podn* |
| 14 | *Lyve1* | *1810032O08Rik* |
| 15 | *Tnn* | *Adam22* |
| 16 | *Tnfrsf9* | *Sgce* |
| 17 | *Nr4a3* | *Cmss1* |
| 18 | *Uck2* | *Tcap* |
| 19 | *Ddit4* | *Gpr153* |
| 20 | *Cxcl1* | *Gm15564* |
| 21 | *Cry1* | *Ckm* |
| 22 | *Tnfrsf21* | *Srrm2* |
| 23 | *Ctsg* | *Car3* |
| 24 | *Slpi* | *Des* |
| 25 | *Lum* | *Rnf207* |
| 26 | *Sncg* | *Acsbg1* |
| 27 | *Pdia4* | *Ckm* |
| 28 | *Mrgprb1* | *Eln* |
| 29 | *Ttyh1* | *Myl1* |
| 30 | *Tpmt* | *Mybpc2* |
| 31 | *Sectm1b* | *Pdlim3* |
| 32 | *Prf1* | *Sfrp2* |
| 33 | *Gzmd* | *Cilp* |
| 34 | *Rpsa-ps10* | *Cygb* |
| 35 | *Mrgprb2* | *Nrap* |
| 36 | *1700071M16Rik* | *Cit* |
| 37 | *Scara5* | *Gm12563* |
| 38 | *Ptgs2* | *Mfap4* |
| 39 | *Notum* | *Gm15662* |
| 40 | *Gzmf* | *Pag1* |
| 41 | *Ccl9* | *Cspg4* |
| 42 | *Mmp3* | *Pdzrn3* |
| 43 | *Adgrf1* | *Cisd3* |
| 44 | *1110038B12Rik* | *Hhipl1* |
| 45 | *Ggt1* | *Flnc* |
| 46 | *Mmp13* | *Cdo1* |
| 47 | *Nr4a1* | *Cacna1g* |
| 48 | *Ppp1r10* | *Xirp2* |
| 49 | *Gzmg* | *Neb* |
| 50 | *Cstf2* | *Cpxm1* |

**Supplementary Table 2:** Top 50 upregulated and downregulated genes from RNA-Sequencing analysis comparing Kin1 WT and Kin1 KO tumours.

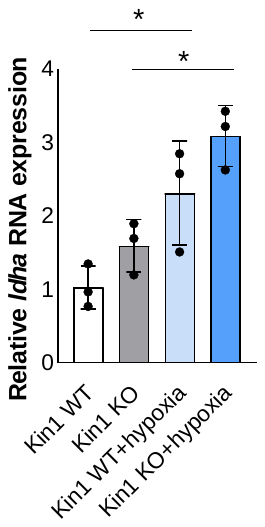

#### Figures

**Supplementary Figure 1: | Confirmation of induced hypoxia in cSCC 2D cultures *in vitro*.** mRNA expression of hypoxic marker *Ldha* at normal and hypoxic conditions in Kin1 WT and Kin1 KO cSCC cells. Data obtained from three independent experiments (mean ± SD). p-values were obtained from one-way ANOVA test followed by Tukey post-hoc test; *p < 0.05.

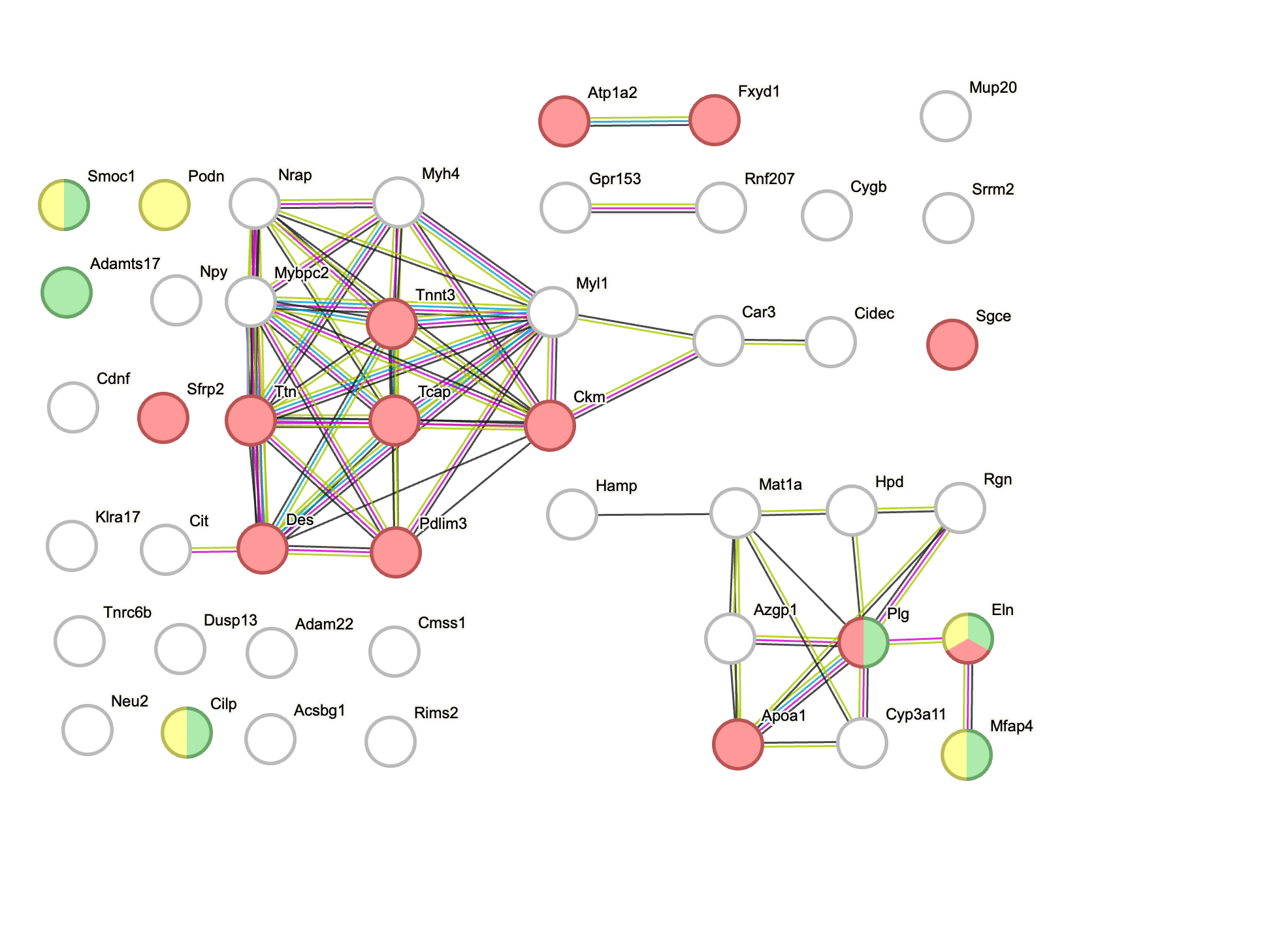

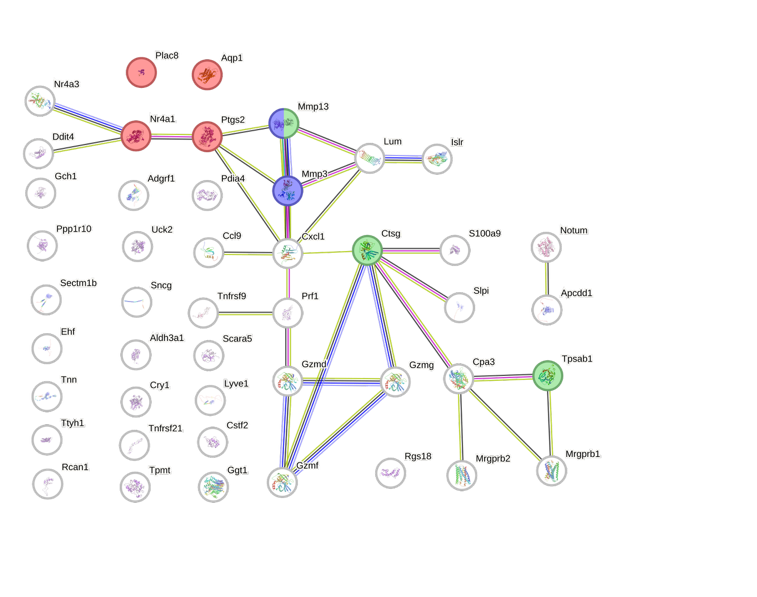

Downregulated genes in Kin1 KO

Upregulated genes in Kin1 KO

**a**

**b**

Extracellular matrix disassembly

Metabolism of angiotensins

Abnormal vascular permeability

Collagen-containing extracellular matrix

Extracellular matrix

Abnormal cardiovascular system physiology

**Supplementary Figure 2 | Differentially expressed genes in Kin1 KO tumours linked to tumourigenic features.** STRING analysis of upregulated (a) and downregulated (b) genes from RNA-Sequencing analysis comparing Kin1 WT and Kin1 KO tumours (n= 3 tumours).

**b** Collagen volume (3D) vs inter-bifurcation λ (3D)

**a** Hypoxic fraction vs vascular density (3D)

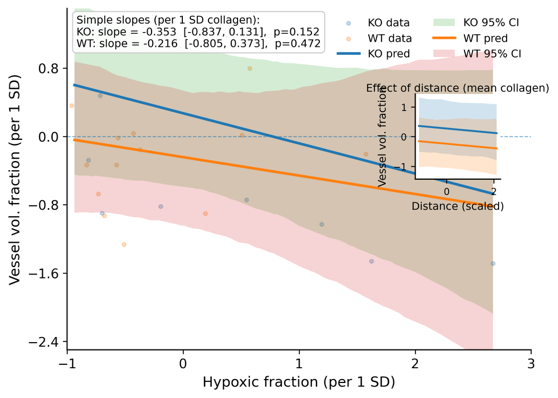

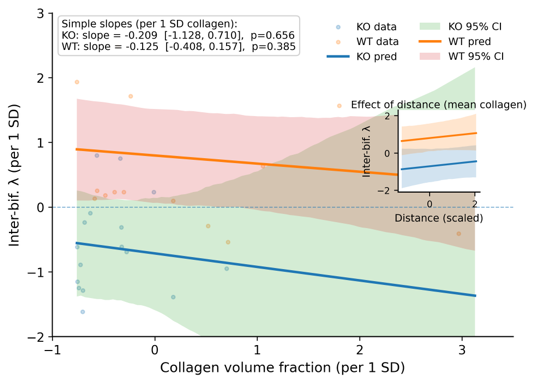

**Supplementary Figure 3 | Further relationships between vascular features, hypoxia and collagen in Kin1 WT and Kin1 KO tumours.** (a) Vessel volume fraction from 3D vessel reconstructions vs hypoxic fraction from 2D cross-sections of the same region, (b) Vascular inter-bifurcation λ (the ratio of vessel length-to-diameter) from 3D reconstructions vs collagen volume fraction, also from 3D. Solid lines represent genotype-specific regressions (KO, blue; WT, orange) with shaded 95% confidence intervals. Insets show the effect of radial distance, confirming independence from spatial gradients.
